# Towards multi-drug adaptive therapy

**DOI:** 10.1101/476507

**Authors:** Jeffrey West, Li You, Jingsong Zhang, Robert A. Gatenby, Joel Brown, Paul K. Newton, Alexander R. A. Anderson

## Abstract

A new ecologically inspired paradigm in cancer treatment known as “adaptive therapy” capitalizes on competitive interactions between drug-sensitive and drug-resistant subclones. The goal of adaptive therapy is to maintain a controllable stable tumor burden by allowing a significant population of treatment sensitive cells to survive. These, in turn, suppress proliferation of the less fit resistant populations. However, there remain several open challenges in designing adaptive therapies, particularly in extending these therapeutic concepts to multiple treatments. We present a cancer treatment case study (metastatic castrate resistant prostate cancer) as a point of departure to illustrate three novel concepts to aid the design of multi-drug adaptive therapies. First, frequency-dependent “cycles” of tumor evolution can trap tumor evolution in a periodic, controllable loop. Second, the availability and selection of treatments may limit the evolutionary “absorbing region” reachable by the tumor. Third, the velocity of evolution significantly influences the optimal timing of drug sequences.

---

Dobzhansky’s now-famous quote that “nothing in biology makes sense except in the light of evolution” succinctly explains a worldview that has been widely adopted by the cancer biology community^1^. Taken one step further, others have claimed that “nothing in evolution makes sense except in the light of ecology” which provided the basis for designing adaptive cancer therapies centered on principles from evolution and ecology^2–4^.

Cancer is an evolutionary and ecological process^5, 6^ driven by random mutations^7, 8^ responsible for the genetic diversity and heterogeneity that typically arises via waves of clonal and subclonal expansions^9, 10^. Clones and subclones compete and Darwinian selection favors highly proliferative cell phenotypes, which in turn drive rapid tumor growth^5, 6^.

Recent emphasis on personalized medicine has largely focused on the development of therapies that target specific mutations. These targeted therapies do extend patient lives but cancer cells tend to evolve resistance within months or years^11, 12^. Prior to therapy, pre-existing resistant cell types are suppressed and kept in check by competitively superior, therapy-sensitive cell types. There is some evidence of “cost” to incurring resistant mutations. In one study, cells sensitive (MCF7) and resistant (MCF7Dox) to doxorubicin cocultured in vitro showed that sensitive MCF7 cells rapidly outcompeted the resistant MCF7Dox line after only a few generations, illustrating the “cost” of resistance cell lines co-cultured with drug-sensitive cell lines.^13^.

With a targeted therapy suppressing sensitive cells, these resistant cell types may experience release from competition^14^. If total eradication of all cancer cells is not accomplished, the tumor will relapse derived from resistant cells that survived initial therapy^15, 16^. Upon relapse, a second drug may be administered. Yet continuous use of this subsequent targeted therapy may inevitably result in the emergence of the corresponding resistant clones (Fig. 1A). This approach ignores considerations of heterogeneity and therapy as a selection event in somatic evolution^9, 17, 18^.

**Figure 1.**
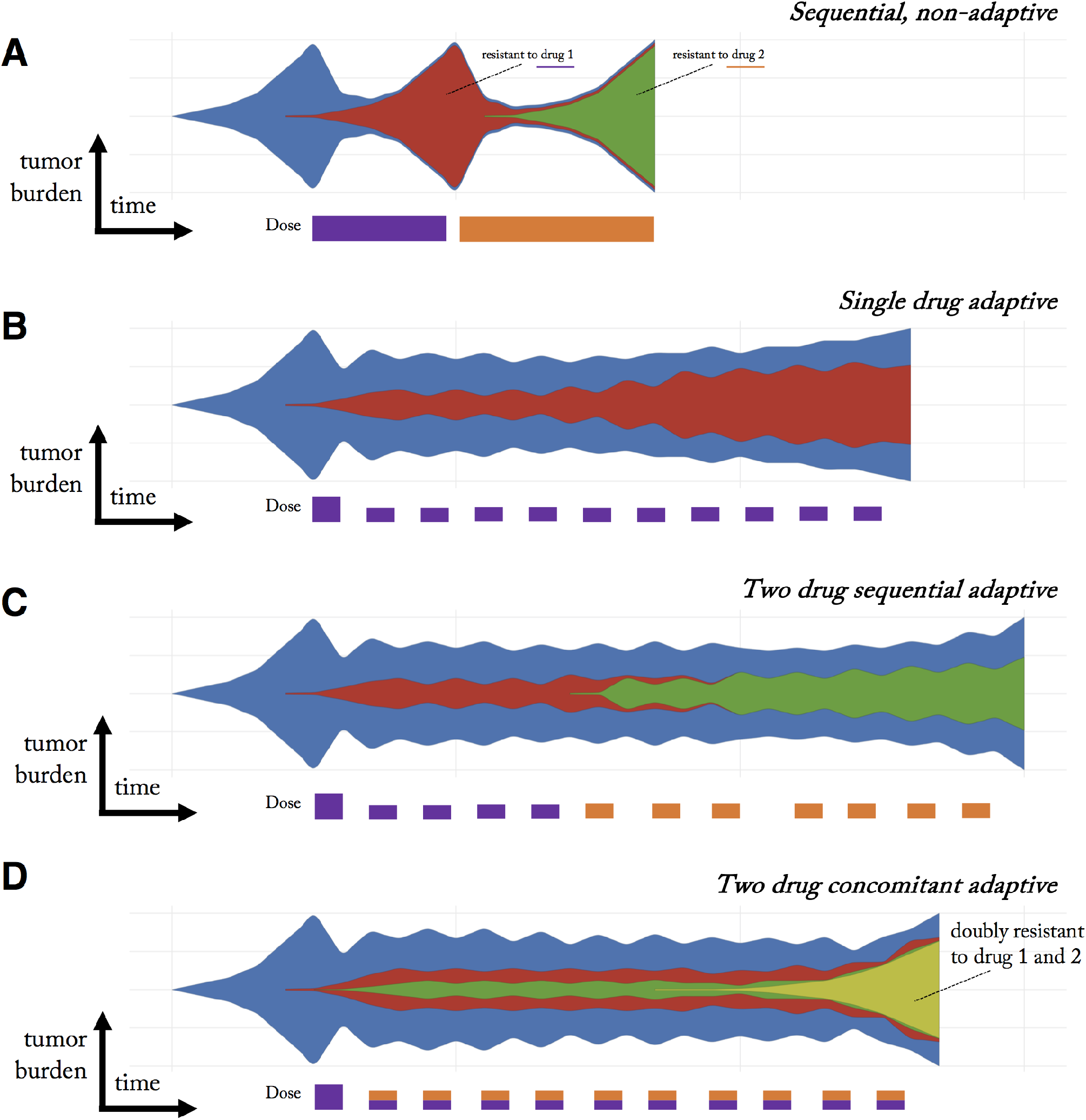
Schematic of cancer clonal evolution under therapy. A) Conventional sequential therapy of two treatments selects for a clone resistant to treatment one (red) upon tumor relapse and subsequently a clone resistant to treatment two (green). B) Adaptive therapy maintains a stable tumor volume by introducing treatment holidays. Drug sensitive clones (blue) suppress the growth of less fit resistant clones (red). However, resistance still eventually occurs. C) One proposed adaptive multi-drug strategy is to alternately switch between drugs during each on-off cycle of tumor burden. D) An alternative multi-drug adaptive strategy is to administer both drugs simultaneously during each on-off cycle, leading to a doubly-resistant resistant clone (yellow).

Historically, the problem of drug resistance has often enlisted the help of mathematical modeling in designing optimal therapy schedules. For example, Goldie and Coldman were the first to propose a mathematical model relating drug sensitivity of tumors to their mutation rates^19^. The model had two clinical implications: first, smaller tumors are more likely to be curable (without pre-existing resistance) and second, as many effective drugs as possible should be applied as soon as possible. They suggest a strict alternating sequence, as many combinations of drugs due to overlapping toxicity and competitive interference. These findings have been nuanced and extended to asymmetrical parameterizations with some success (see e.g.^20^). In this manuscript, we will explore sequential and concomittant therapy, with the goal of designing patient-specific therapeutic schedules which drive tumor evolution into cycles, explained below.

## Enlightenment via evolution

Eradicating most disseminated cancers may be impossible, undermining the typical treatment goal of killing as many tumor cells as possible^21, 22^. Previous schools of thought saw maximum cell-kill as either leading to a cure or, at worst, maximizing life extension. Attempting to kill as many cancer cells as quickly as possible may be evolutionarily unsound and facilitate resistance evolution and loss of therapy efficacy. Evolution matters. A recent (2012) systematic literature analysis of cancer relapse and therapeutic research showed that while evolutionary terms rarely appeared in papers studying therapeutic relapse before 1980 (< 1%), the use of evolutionary terms has steadily increased more recently, due to the potential benefits of studying therapeutic relapse from an evolutionary perspective^23^.

A new paradigm in the war on cancer replaces the “treatment-for-cure” strategy with “treatment-for-contain” – receiving cues from agriculturists who have similarly abandoned the goal of complete eradication of pests in favor of more limited and strategic application of insecticides for control^21, 24^. This eco-evolutionary inspired paradigm for cancer treatment known as “adaptive therapy” capitalizes on subclonal competitive interactions. Resistance may confer some fitness costs due to increased rates of DNA repair or other costly activities required to pump toxic drugs across cell membranes. Cancer cell resistance mechanisms, whether mitigation, detoxification, or re-routing metabolic pathways, divert finite resources that would otherwise be available for cell proliferation or other avenues for cell survival^14, 22, 25^.

The goal of adaptive therapy is to maintain a controllable stable tumor burden by allowing a significant population of treatment sensitive cells to survive (see Fig. 1B, blue). These readily treatable sensitive cell serve to supress the proliferation of the less fit resistant populations (see Fig. 1B, red). Adaptive therapies have now been tested experimentally^26, 26, 27^ and are currently being applied across multiple clinical trials (NCT02415621; NCT03511196; NCT03543969; NCT03630120) at the Moffitt Cancer Center^28^. These adaptive therapies capitalize on competition for space and resources between drug-sensitive and slow growing drug-resistant populations^13, 29, 30^. These trials have shown initial promise in prostate cancer compared with contemporaneous patient cohorts^28^, but it should be noted that evolutionary modeling-based dosing schedule did not improve progression-free survival in anti-cancer TKI regimens for patients with EGFR-mutant lung cancers^31^.

## Steering patient-specific evolution

This adaptive approach means that each patient’s treatment is truly personalized based on the tumor’s state and response rather than a one-size-fits-all fixed treatment regime^32^. There remain several open challenges in designing adaptive therapies. First, treatments can aim to steer and control the eco-evolutionary dynamics to where the tumor finds itself in an evolutionary dead end^33^, an evolutionary double-bind^34–36^, or evolutionarily stable control^13, 21, 37^. Yet, for clinical practice, how to design such therapies remains difficult^38^. Second, it’s not yet clear how to extend these evolutionarily enlightened therapeutic concepts to multiple treatments. Two schematic examples are illustrated in Fig. 1C and 1D. Is it evolutionarily optimal to reproduce the single drug adaptive therapy in a sequential (Fig. 1C) setting or in concomitant (Fig. 1D)? Synergizing treatments such that evolving resistance to one drug makes cells more susceptible to another requires mathematical modeling as well as improved monitoring methods^39, 40^.

Figure 2 provides a schematic of steering the tumor into “cycles” of tumor evolution. A cycle is defined as a treatment regimen that steers the tumor into periodic and repeatable temporal dynamics of tumor composition. As seen in figure 2, a weekly biopsy shows the frequencies (pie charts) of 4 phenotypes (blue, green, red, yellow) in the evolving tumor over time. At each time step, the state of the tumor is given by a vector, 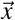, indicating the frequency of each cell type composing the tumor. In this example, the fifth week produces a tumor composition, 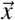, which is approximately equivalent to the tumor composition at the start of therapy 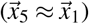. Theoretically, such cycles could be repeated ad infinitum to steer and trap tumor evolution in a repeatable (and controllable) cycle.

**Figure 2.**
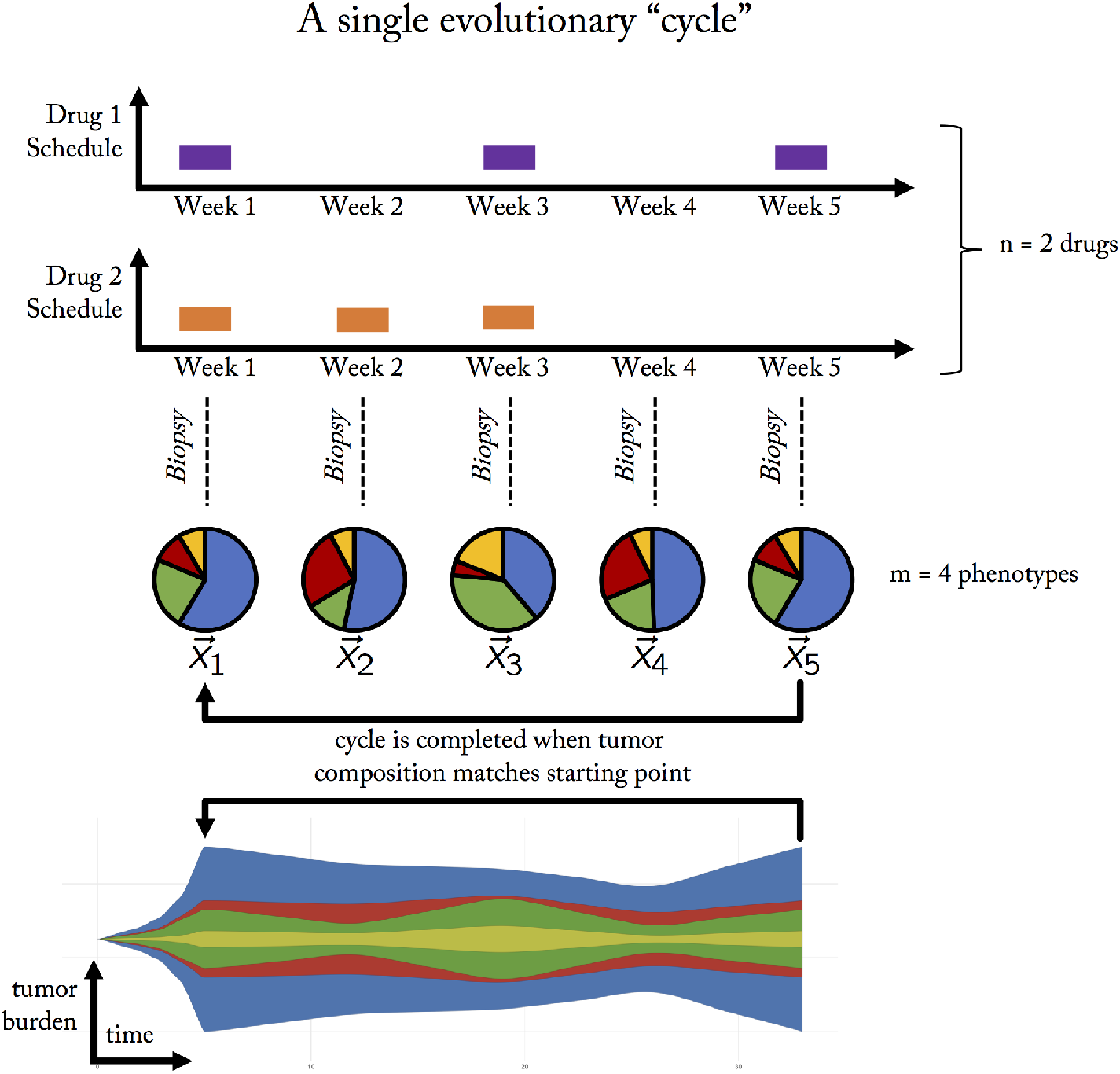
A single evolutionary cycle. Two drugs (purple, orange) are administered either in combination (e.g. week 1), monotherapy (week 2, 5) or no treatment (week 4). A tumor biopsy indicates the tumor composition at the end of each week, showing the evolution of four phenotypes (pie charts: blue, green, red, yellow phenotypes) over time. The goal of adaptive therapy is to maintain a stable tumor volume while controlling the composition of the tumor with respect to cell type frequencies (stored in the state vector, 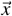). A frequency-dependent “cycle” is a paradigm of tumor control which employs a succession of treatments that returns the state of the tumor composition back to the initial state. A cycle of five weeks is shown here: 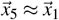. In this example, it is assumed that a single unvarying course of action would result in an unacceptable tumor composition of resistant cell types.

A trial may use sequences of *n* available drugs either alone or in combination (shown for *n* = 2 in figure 2). We employ the terminology “treatment” to indicate the 2^*n*^ possible combinations: no drug, single drug, or combination therapy. These treatments can be administered in any arbitrary sequence with the goal of controlling *m* cell types. An adaptive trial for metastatic castrate resistant prostate cancer (NCT02415621) uses only two treatments: 1. Lupron and 2. Lupron & Abiraterone. Likewise, an adaptive trial for advanced BRAF mutant melanoma (NCT03543969) administers Vemurafenib and Cobimetinib in combination, followed by no treatment. Both trials use only two treatments out of the four (2^2^) combinations possible with two drugs (no treatment; first drug only; second drug only; first and second drug in combination therapy). Opening up trial design to include the full range of complexity (i.e. 2^*n*^) may allow for greater tumor control, but the treatment administered at each clinical decision time point must be chosen with care and forethought, to steer the tumor into a desirable evolutionary state.

Adaptive therapy has been attempted clinically using proxy measurements for tumor volume such as prostate-specific antigen marker (PSA) for prostate or lactate dehydrogenase (LDH) for melanoma. While these blood biomarkers are crude measurements of tumor burden, there have been many recent advances in reliable methods of monitoring tumor evolution using circulating tumor DNA as an informative, inherently specific, and highly sensitive biomarker of metastatic cancers (see^41^). As monitoring methods mature, so must methods of prediction and quantification of evolution^28, 42, 43^. To that end, the purpose of this paper is to introduce the following three concepts to consider in the pursuit of designing multi-drug adaptive therapies:

1. Frequency-dependent “cycles” of tumor evolution
2. Treatment-dependent evolutionary “absorbing region”
3. Frequency-dependent evolutionary velocities

A significant issue however, is that it is unclear when such cycles occur and how to choose between multiple cycling options. To that end, we propose the concepts of “absorbing regions” and “evolutionary velocities” to aid the exploration and selection of adaptive evolutionary treatment cycles. This adaptive approach to therapy maintains tumor volume in favor of controlling the tumor composition (i.e. mitigating the onset competitive release of resistance).

## Results

In progressive prostate cancer, continuous or intermittent androgen deprivation (IAD) are common treatments that generally leads to a substantial decline in tumor burden. Eventually, tumor burden rebounds as a result of the rise of castrate-resistant cells^44^. Upon castrate-resistance, adaptive therapy using abiraterone has shown considerable treatment success. Previously, this disease was described by a population dynamics model with three cell types: those that require testosterone for growth, *T*^+^, cells that require testosterone but produce their own, *T^P^*, or cells that are independent of testosterone, *T*^−28^. This model served to motivate a clinical trial (NCT02415621) which administers continuous Lupron (limiting systemic testosterone production) and cycles of adaptive Abiraterone (anti-androgen). This model suggests that even with adaptive therapy, disease progression can be greatly delayed but remains inevitable (see figure 3A,C). In figure 3, we present data of representative patients from NCT02415621 who have undergone at least five adaptive doses of treatment at the time data was acquired^28, 38^. Next, we parameterize a previously published Lotka-Volterra model of treatment dynamics (see refs.^28, 38, 45^), by fitting the model to prostate-serum-antigen blood biomarker (PSA) data from all patients (figure 3, dashed red lines).

In order to demonstrate the feasibility of our proposed evolutionary cycling concept, a new adaptive treatment schedule is proposed (figure 3, blue line) for each patient. The treatment schedule is constructed such that the tumor subpopulations return to the original proportion and thus complete, as proposed in the above figure 2. Three treatments are given: no treatment (purple), Lupron only (blue), Lupron & Abiraterone (red), shown for twenty repeated cycles. This patient-specific cycle which prolongs the relapse of the resistant *T*^−^ subpopulation while maintaining a controllable tumor volume.

### Case study: metastatic castrate resistant prostate cancer

Figure 3 indicates the promise of designing treatment schedules to drive tumor evolution into a cycle, but it is unclear when such cycles occur. To help elucidate the nature of such cycles, we simplify our model system by ignoring (for the moment) population dynamics to leverage several mathematical conveniences of a frequency-dependent game theoretic modelling framework, explained below. Frequency-dependent models of tumor evolution are particularly suited for studying tumor control (i.e. “cycles” of tumor evolution), determining the set of possible evolutionary dynamics (i.e. evolutionary absorbing region), and timing of evolution (i.e. evolutionary velocity).

This case study introduces and develops these generalizable concepts for designing multi-drug adaptive therapy treatment schedules for three cell types (m = 3 phenotypes; figure 2) under two drug treatments (n = 2 drugs; figure 2). In principle, the concepts applied to this particular case study can be extended to an arbitrary number of available treatment combinations for desired control of an arbitrary number of cell types.

In order to track the state vector of the tumor, 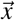, we set the variables *x*_1_, *x*_2_, *x*_3_ to the corresponding frequency of dependent (*T*^+^), producers (*T^P^*) and independent (*T*^−^) cells, respectively, such that 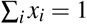. The temporal dynamics of these different populations under treatment can be characterized inside a trilinear coordinates simplex (Fig. 4A) which gives a representation of every possible value of 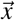, on a triangle. The corners represent a tumor consisting of 100% of either *T*^+^ (top corner), *T^P^* (left corner), and *T*^−^ (right corner). Figure 4 shows the temporal dynamics under each treatment.

Every treatment scenario has an associated long-time equilibrium state: the tumor composition where all temporal dynamics eventually converge. For example, continuous treatment of Lupron leads to roughly equal fraction of *T*^+^ and *T^P^* (blue circle Fig. 4B). Continuous treatment of Lupron & Abiraterone leads to full saturation of *T*^−^ (blue circle Fig. 4C). Similarly, administering no treatment leads to an equilibrium of mostly *T*^+^ and roughly equal (but small) proportions of *T*^−^ and *T^P^* (purple circle Fig. 4D).

**Figure 3.**
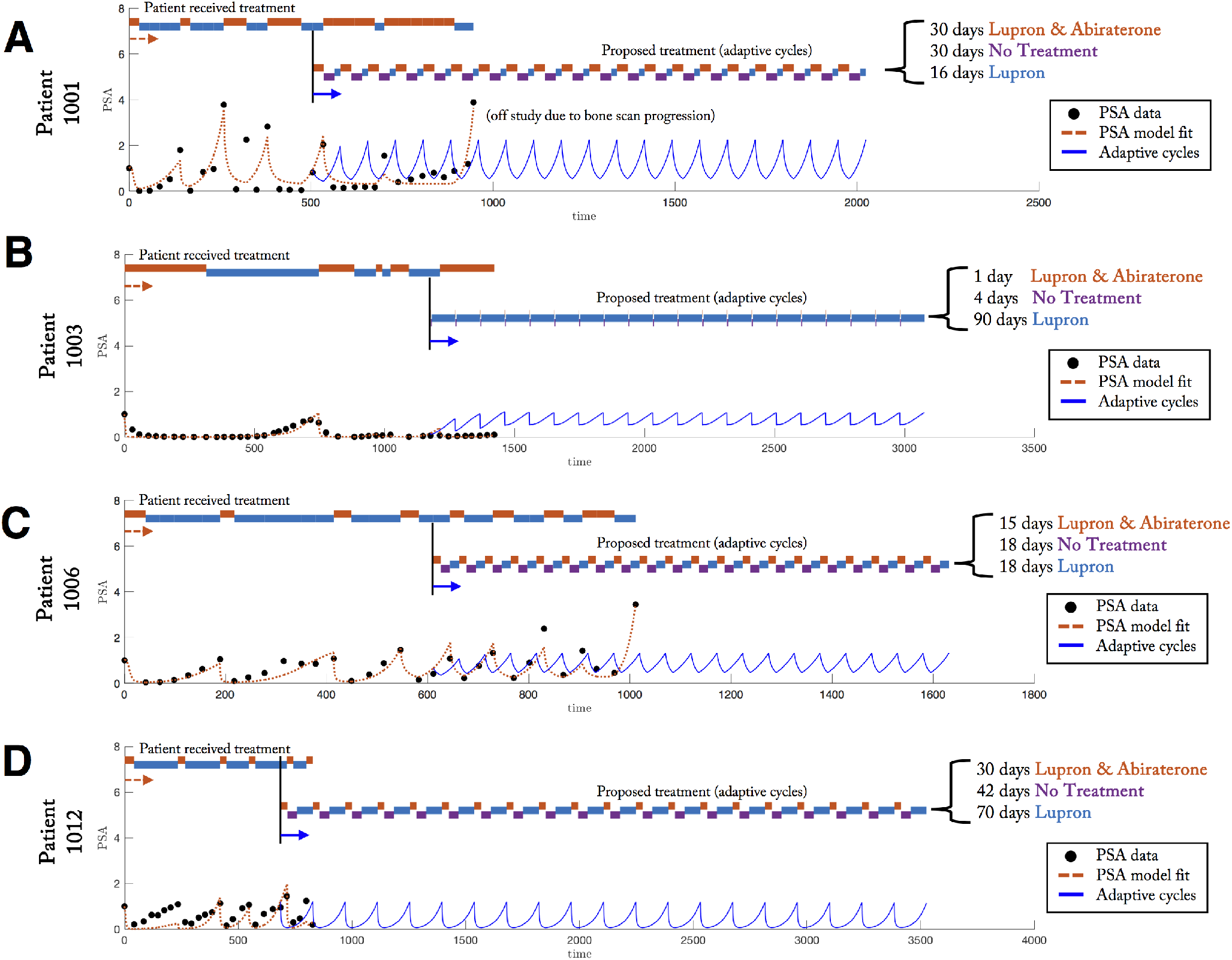
Patient-specific evolutionary cycles delay emergence of resistance. Model fit to PSA data from clinical trial NCT02415621, which administers continuous Lupron with adaptively administered Abiraterone, is shown (black circles). A previously published model is fit to the data (see^28, 38^, red dashed line). Timing of treatment received under clinical protocol is indicated (top) alongside a proposed treatment of “adaptive cycles” (model-predicted PSA shown in blue). This adaptive cycling approach is able to control the tumor (shown for twenty cycles).

**Figure 4.**
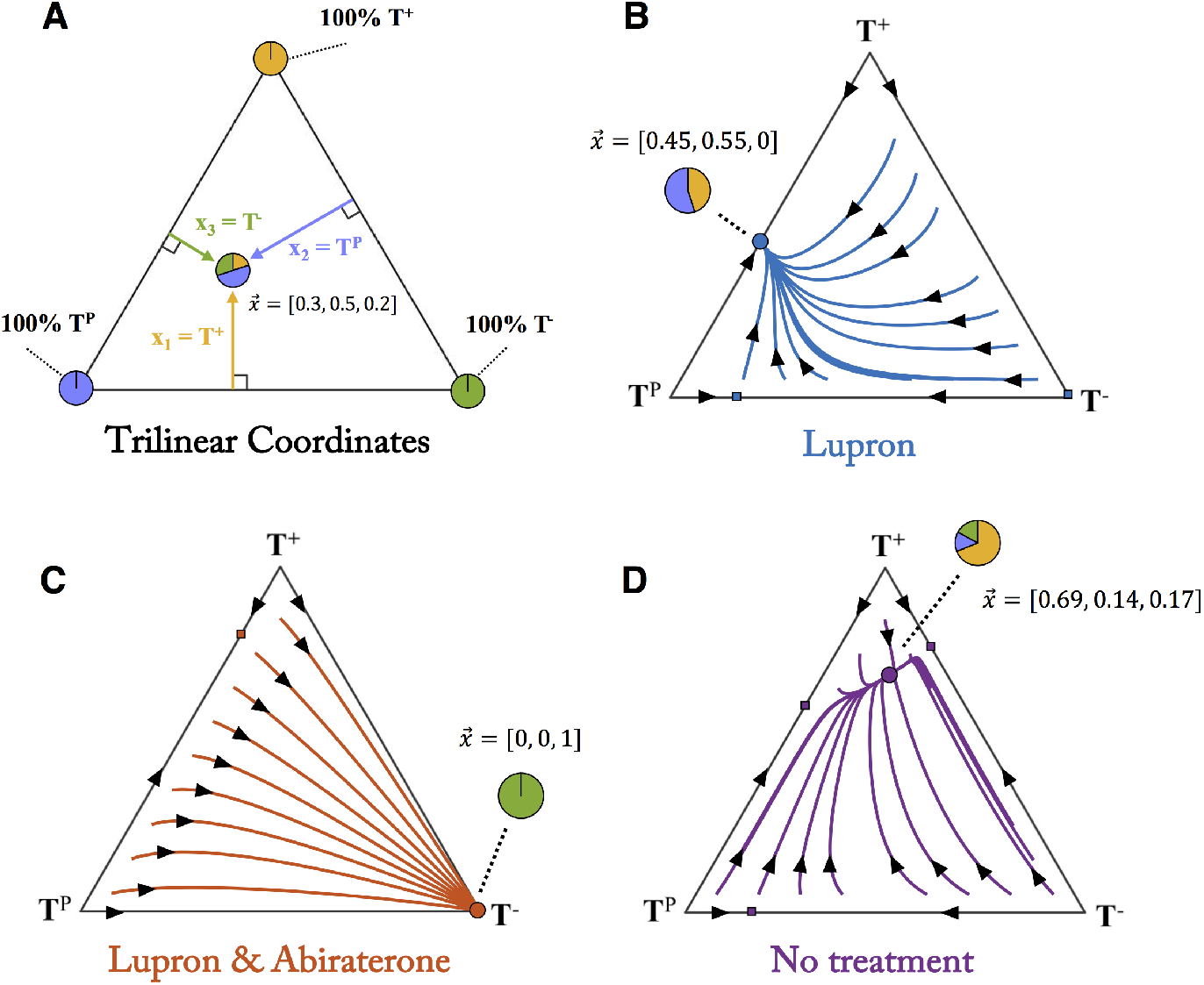
Temporal dynamics under therapy. (A) A “trilinear” coordinate system represents every possible state of the tumor’s 3 cell types 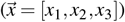 inside the triangle. The corners represent 100% of each particular cell type while interior points represent a fraction of each cell type. (B) Temporal dynamics are traced along trajectories for Lupron treatment (blue lines with arrows). Each point inside the triangle leads to an equilibrium state of *T^P^* and *T*^+^ cells (blue circle). (C) Similarly, Lupron & Abiraterone treatment leads all dynamics to the 100% *T*^−^ equilibrium state (red circle). (D) The tumor still evolves without treatment, to an equilibrium of mostly *T*^+^ cells (purple circle).

This leads us to a key insight. We are able to ignore the tumor volume information (for the moment) because of one convenient fact: an optimal treatment schedule will necessarily be a schedule which successfully avoids all equilibria. Equilibria associated with “no treatment” will lead to tumor saturation and presumably, death. Equilibria associated with any treatment will lead to a fully resistant tumor (and also presumably, death). For the moment, we ignore tumor volume in favor of designing schedules that steer evolution into adaptive cycles, with our implicit assumption that cycles far from any associated equilibria will outperform cycles near equilibria.

The goal of adaptive therapy in prostate cancer is to delay the onset of the resistant *T*^−^ population by well-timed *switching* between each treatment. This is equivalent to switching between each triangular phase portrait in figure 4 before reaching the resistant equilibrium state of any given treatment. In theory, a periodic (closed) cycle can be constructed by switching between treatments at carefully chosen times in order to design schedules that are superior to continuous maximum tolerated dose^46^.

### Searching for cycles

For example, figure 5A shows a patient with an initial condition of roughly equal fractions of each cell type (blue circle). Lupron might be administered for some time (blue line) before switching to no treatment (purple line), followed by a switch to both Lupron & Abiraterone (red line). This sequence of treatments arrives back at the same initial condition: an evolutionary cycle. This treatment sequence can be repeated, controlling the tumor. The dashed lines on figure 5A show dynamics under continuous treatment which lead to respective equilibrium states, spiralling out of the evolutionary cycle.

**Figure 5.**
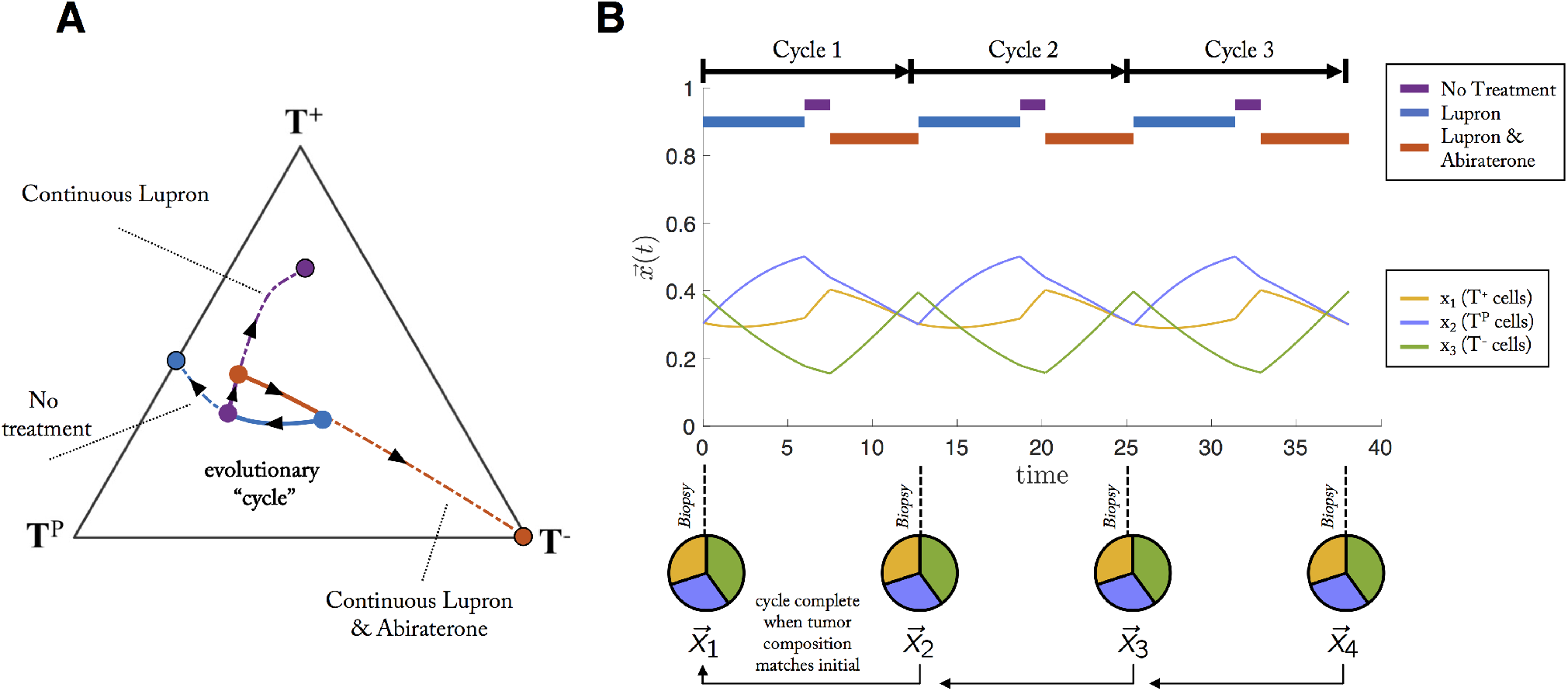
Sequential treatments which lead to an evolutionary cycle. An evolutionary cycle employs a succession of sequential treatments which returns the state of the tumor composition back to the initial state at the start of treatment. (A) For example, a patient with an initial condition of roughly equal fractions of each cell type (blue circle) might be administered Lupron for some time (blue line), switch to no treatment (purple line), and switch to Lupron & Abiraterone (red line), which arrives back at the same initial condition: an evolutionary cycle. This progression can be repeated, controlling the tumor. Alternatively, the dashed line show dynamics under continuous treatment, leading to respective equilibrium states, spiralling out of the evolutionary cycle. (B) The identical evolutionary cycle paradigm is shown with cell fraction, 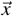 over time.

The identical evolutionary cycle paradigm is shown with cell fraction, 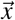 over time in figure 5B. By appropriate treatment switching, the three cell types remain in competition with each other, and no cell type is able to dominate but instead are balanced indefinitely in closed periodic cycles thereby avoiding the emergence of resistance. The tumor undergoes 3 such evolutionary cycles, where the tumor “resets” back the initial state before treatment (pie charts, figure 5).

From this perspective, adaptive therapy may benefit from treatment holidays precisely because these holidays help the tumor composition be pushed into an approximate cycle. Yet, when slightly off, resistance still *eventually* occurs in adaptive therapy regimes when not on an exact cycle (Fig. 1B, red). By observation, many such cycles exist in the state space shown in Fig. 4D: for example, traveling down any Lupron trajectory (blue), switching to a no treatment trajectory (purple) and so on, repeated ad infinitum. Where do such cycles exist? Which cycles are preferable? To answer these questions, the next section introduces concepts of evolutionary absorbing region and evolutionary velocity.

### Evolutionary absorbing region

These same phase portraits can be drawn for each pairwise treatment combination (Fig. 6). Each treatment has an associated equilibrium (solid circles), which can be connected by a single evolutionary trajectory. These two connecting trajectories represent a bounding region of state space where a periodic cycle resulting from sequential administration of the two treatments is guaranteed on the outer rim of this region (directionality shown with black arrows). As can be seen in Fig. 6, all external trajectories tend toward either treatment’s equilibrium, or towards the inside of the bounding domain. The implications are clear: sequential treatment between any two drugs for a sufficiently long time results in limiting the “evolutionary absorbing region” of the tumor composition.

**Figure 6.**
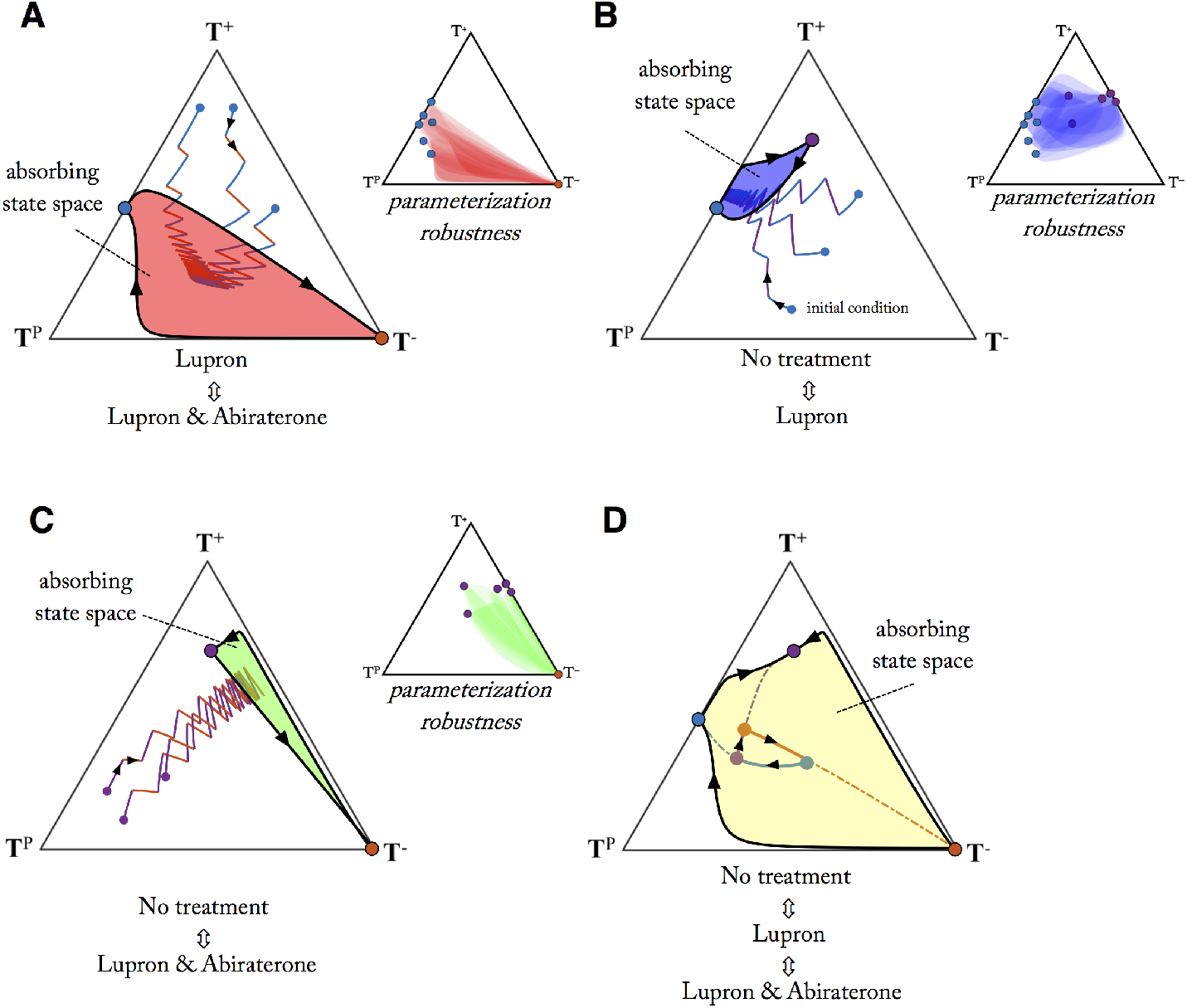
Evolutionary absorbing regions under therapy. Bounded, absorbing regions are shown for treatment pairs: Lupron ↔ Lupron & Abiraterone (A); No treatment ↔ Lupron (B); No treatment ↔ Lupron & Abiraterone, (C) and all treatments (D). The outer rim of the absorbing region represents the largest cycle for the pair of treatments (arrows show trajectory directionality). Any regimen will eventually be absorbed into the bounded region (3 sample sequential regimens shown). By cleverly defining treatment schedules (D), the bounded region can be expanded to find cycles in a wider region of the state space. Example shown for Lupron (blue line) → no treatment (purple line) → Lupron & Abiraterone (red line). Parameter robustness (inset shown in A,B,C): absorbing state space is shown for 36 random parameterizations subject to inequalities discussed in table 1 and 2, overlaid in transparent color (see Methods). For these given inequalities, the absorbing regions remain relatively robust to parameter changes.

Herein lies a second key advantage of frequency-dependent game theoretic models: parameterization robustness. Shown in the inset of each subfigure (figure 6), are 36 random parameterizations (subject to the inequalities in table 1 and 2), overlaid in transparent shading. Absorbing regions overlap significantly due to the fact that model dynamics are more sensitive to *relative* parameter values (i.e. the inequalities in table 1 and 2) than *absolute* values chosen.

**Table 1.** Competition parameters for Lupron and Abiraterone treatment

|  |  |
| --- | --- |
| $c > e$ | $T^P$ cells have a higher fitness than $T^-$ cells when interacting with few $T^+$ (absence of competition) |
| $a > f$ | Interacting with mostly $T^P$ cells, $T^+$ gains from the public good and from the new available space in low vasculature regions |
| $b < d$ | Interacting with mostly $T^-$ cells, $T^P$ cells see little competition near vasculature. Payoffs to $T^+$ cells, $b$, may be small or zero |
| $a > b = 0$ | $T^+$ cells need the $T^P$ cells to succeed in the absence of systemic testosterone |
| $c > d$ | $c$ is likely the largest parameter as $T^P$ cells have the highest fitness in a mostly $T^-$ tumor without systemic testosterone |
| $e > f$ | Again, $T^P$ cells outcompete $T^-$ cells in the absence of systemic testosterone |

**Table 2.** Competition parameters for no treatment

|  |  |
| --- | --- |
| $c < e$ | $T^-$ cells have a higher fitness than $T^P$ cells when interacting with many $T^+$, especially in low vasculature regions. Testosterone production by $T^P$ production comes at some cost to provide public good to both self ($T^P$) and neighbor ($T^+$). Both $c$ and $e$ should decrease in the pre-treatment condition |
| $a > f$ | $T^+$ cells have a higher fitness than $T^-$ cells when interacting with many $T^P$, receiving advantage from the public good. The parameter $a$ should increase in the pre-treatment condition $f$ slightly decrease |
| $b > d$ | $T^+$ cells have a higher fitness than $T^P$ cells when interacting with many $T^-$ because there is lack of spatial competition near vasculature for $T^+$ cells as testosterone is not being used |
| $a < b$ | $T^+$ cells have a higher fitness competing with $T^-$ over competition with $T^P$ |
| $c < d$ | Similarly, $T^P$ cells have a higher fitness competing with $T^-$ over competition with $T^+$ |
| $e > f$ | $T^-$ cells have less competition for space in a tumor with mostly $T^+$ than with mostly $T^P$. The parameter $f$ should decrease in the pre-treatment condition slightly |

At least one cycle exists for each pairwise treatment scenario: the outer rim of the absorbing state. There may be other cycles, existing only in the interior of the region. For some pairwise combinations, the bounding region is quite large (see Fig. 6A: Lupron ↔ Lupron & Abiraterone); for others quite small (see Fig. 6B: no treatment ↔ Lupron or C: no treatment ↔ Lupron & Abiraterone). Once treatment drives the tumor composition into one of these bounding regions, it’s impossible to traverse outside without the addition of a third treatment.

Considering the three treatment scenarios together (see Fig. 6D: no treatment ↔ Lupron ↔ Lupron & Abiraterone) expands the evolutionary absorbing region. Arbitary cycles can be constructed by clever sequences of treatment inside this expanded evolutionary absorbing region. To illustrate which regions in which cycles may be preferable, the next section will describe evolutionary velocity.

### Evolutionary velocity

The evolutionary “velocity” (and each of its velocity components) of each treatment scenario can be viewed on the same trilinear simplex, seen in Fig. 7 for no treatment (7, left column), Lupron (7, middle column), and Lupron & Abiraterone (6, right column). A low velocity indicates a slow change to the tumor composition (calculated as the magnitude of the resultant vector of all components in eqn. 1: see Supporting Information). Several recent studies have indicated the importance of studying temporal effects of drug administration. Using human breast cancer explants, in vitro cells, mouse in vivo studies combined with mathematical modeling, one study showed that a temporary exposure to a taxane induces phenotypic cell state transition towards a favored transient chemotherapy-tolerant state which disappears when the drugs are administered concomitantly^47^. A separate study noted the appearance of weakly proliferative, resistant cells under high dose therapy is transient and controllable by non-genetic, stem-like (reversible) characteristics that depend on timing and length of drug administration^48^.

**Figure 7.**
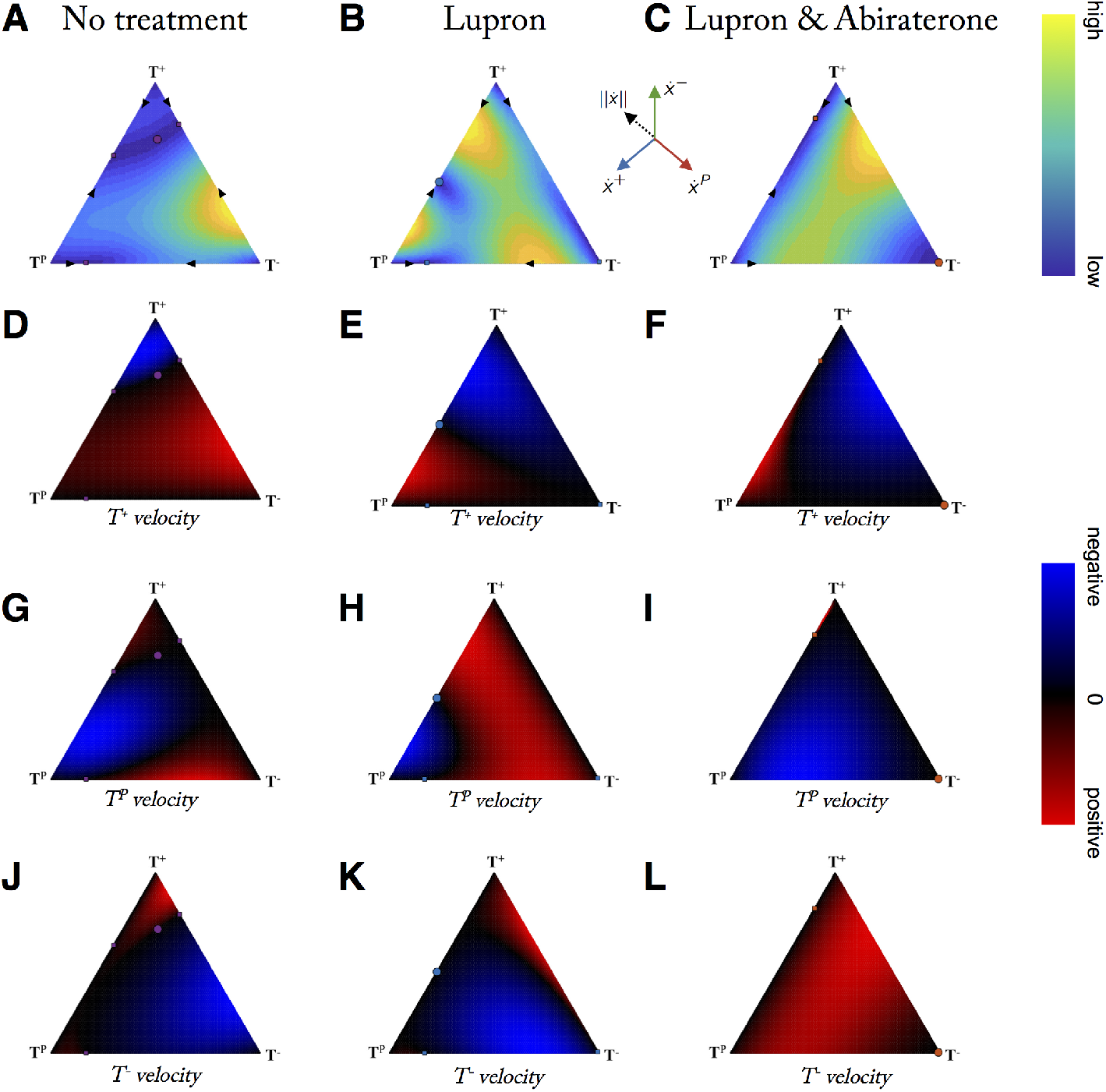
Evolutionary velocity before and during therapy. Columns show evolutionary velocity of simulations identical to Fig. 4 for no treatment, lupron, and lupron+abiraterone. A-C: note that no treatment is characterized by mostly slow velocities (blue) while adding treatments increases velocities, except when approaching evolutionary stable points (circles). D - L: rows show subpopulation relative velocity for T+ (D,E,F), TP (G,H,I), and T− (J,K,L). Adaptive therapy relies on the suppression of the resistant population (T-) by the sensitive populations (T+,TP) during treatment holidays (first column). This is only possible in certain a subset of the statespace (blue; where T− velocity < 0). Velocities calculated by eqn. 1: see Supporting Information.

In short: timing matters. Introduction of concomitant therapy or altering the timing of sequential therapy changes the evolutionary velocity of the underlying tumor composition. One can imagine scenarios where a fast or slow evolutionary velocity is desired. A tumor with a high composition of resistant cells may need to navigate to a fast dynamics region to rapidly decrease the resistant subpopulation. Alternatively, an adaptive regime may capitalize on slow velocities regions that ensure slow evolutionary dynamics on treatment holidays.

Frequency dynamics models allow for monitoring and control of the velocity of a single cell type (see Fig. 7 D - L). For example, under continuous Lupron & Abiraterone treatment, the velocity of the *T*^−^ population is positive for most of the state space (see Fig. 7L). In order to control the *T*^−^ population, it’s necessary to switch to a new treatment: Fig. 7J or Fig. 7K. Depending on the current tumor composition, no treatment or Lupron may be desirable for negative *T*^−^ velocity (blue). For example, while certain adaptive evolutionary cycles may be *technically* feasible, clinical considerations of fast evolutionary velocities may preclude certain schedules requiring high frequency hospital visits or impractically slow wash out times of particular treatments.

## Discussion

In the design of multi-drug adaptive therapy, the optimal method of combining therapies in an additive or sequential manner is unclear. A key observation is that the goal of adaptive therapy is to maintain a stable tumor volume in favor of designing therapies that alter tumor composition. We advocate for the use of frequency-dependent competition models (and in particular evolutionary game theory) to design novel multi-drug adaptive therapy regimens. This is a promising modeling approach, and more work must be done combining population models of changing tumor volume with frequency dependent models of changing tumor fraction in order to eliminate clinically unfeasible treatment schedules from consideration.

While it’s clear that tumors evolve in response to treatment, it’s proved difficult to exploit evolutionary principles to steer tumor evolution in an ideal direction. One limitation of current adaptive therapy clinical trials is the limited selection of drugs and limited monitoring methods. Even despite the lack of monitoring the exact state of the tumor composition, the technique of maintaining a substantial drug-sensitive population for extended tumor control has shown promise in mathematical models as well as preclinical and clinical trials. However, each of the on-going or planned adaptive therapy clinical trial utilizes less than the full range of treatment options available. For example, prostate cancer adaptive therapy sequentially administers Lupron and both Lupron & Abiraterone, while ignoring no treatment or Abiraterone monotherapy. Similarly, the planned melanoma trial administers no treatment alternating with both Vemurafenib and Cobimetinib. The conceptual paradigms introduced here (cycles of tumor evolution, evolutionary absorbing region, and evolutionary velocity) provide a path forward to reducing the complexity of selecting between 2^*n*^ treatment choices at every clinical decision point.

Another trial, currently accruing castrate-sensitive prostate cancer patients (NCT03511196), attempts to infer underlying dynamics of tumor subpopulations by comparison two clinical biomarkers over time: PSA and testosterone levels. Patients with rising PSA levels without a respective rise in testosterone may indicate an emergent castrate-resistant population (TP). When this occurs, abiraterone is adminsitered to counter this resistant population while leaving the serum testosterone unchanged to bolster the T+ population. Even without precise monitoring of underlying subpopulations, changes may be inferred using evolutionary principles.

Especially when developed in collaboration with clinicians, evolutionary models of cellular competition can provide clarity and power despite their simplicity. As a first step, we have illustrated these three adaptive paradigms in a specific case study (prostate cancer) for *n* = 2 drugs controlling *m* = 3 phenotypes (testosterone producers, testosterone dependent, testosterone independent cells). These techniques can be extended to any multi-drug adaptive therapy setting. The first step is to carefully choose competition parameters (i.e. the payoff matrix) of each cell phenotype under consideration and draw the dynamical phase portraits of possible evolutionary trajectories implies by the set of competition parameters. The power of evolutionary game theory is that the *relative* fitness of each phenotype is often easy to determine, allowing for straightforward construction and analysis of the model.

Each treatment will eventually fail due to resistance. The resistant state is the absorbing stable state of tumor phenotypic composition associated with continuous treatment. Example trajectories for a given treatment can be plotted, predicting the timing to reach this absorbing resistant state. When adding a second treatment in sequence, an absorbing bounded region of phenotype space (Fig. 6) can be drawn, which we term the evolutionary “absorbing region” reachable with two or more treatments. This absorbing state space is bounded by the evolutionary cycle that connects the stable state points from each treatment (note: we consider points in the interior of the simplex and the global equilibrium only). The absorbing region may be be a small region (i.e. Fig. 6B) or large (i.e. Fig. 6A). Upon combining all treatments, the state space is expanded, also leading to more options for cycles in new regions of the state space. Intuition would indicate that new drugs should be introduced that have “orthogonal” stable states – stable equilibria far from the equilibria of existing drugs. These orthogonal drugs open up the treatment space, allowing for more options in choosing cycles (Fig. 6D). The closeness of two stable states from two distinct treatments will give an idea of the “orthogonality” of two treatments, aiding treatment selection, the sequential ordering of treatments, and the timing of switching between treatments.

It is also important to note that not treating still allows the tumor to evolve. This no treatment case is often associated with slow evolution due to low selection. During treatment holidays used to avoid resistance to a first treatment, it may be wise to choose a faster (high selection) second treatment that gives a similar resultant tumor composition over the slow evolution of no treatment. Adaptive therapy is a promising step towards ecologically-inspired personalized medicine. Optimizing multi-drug adaptive therapy is not a straightforward task, but mathematical models are a powerful abstraction of clinical intuition, enabling the generation of new treatment schedules and comparisons to standard of care.

## Methods

### Frequency Dynamics

Evolutionary game theory (EGT) is a mathematical framework that models frequency-dependent selection for strategies (phenotypes) among competing individuals. Competition between individuals is typically characterized by a “payoff matrix” which defines the fitness of an individual based upon interactions with another individual or the population at large^49–51^. As a game, the payoff to an individual depends both on its strategy and the strategies of others in the population. As an *evolutionary* game, the payoffs to individuals possessing a particular strategy influences the changes in that strategy’s frequency. A strategy that receives a higher than population-wide average payoff will increase in frequency at the expense of strategies with lower than average payoffs. Such frequency-dependent mathematical models have shown success in modeling competitive release in cancer treatment^14^, designing optimal cancer treatment^52–54^, evolutionary double binds^35^, glioblastoma progression^55^, tumor-stroma interactions^56^, the emergence of invasiveness in cancer^57^ as well as in co-cultures of alectinib-sensitive and alectinib-resistant non-small cell lung cancer^58^.

In principle, there exist many frequency-dependent models of cell-cell competition which could adequately characterize this system: replicator dynamics models, stochastic Moran process models, spatially-explicit game theoretic representations, even normalized population dynamics models. Here, we simply require a model which analyzes trajectories of *relative* population sizes rather than *absolute* population sizes of *m* cell types under treatment from combinations of *n* drugs. For details on the specific implementation and parameterization of the model used in figures 4, 6, and 7, we refer the reader to Supplemental Information and to refs.^28, 45^.

The model presented below is a simplified frequency-dependent dynamics mathematical model (a qualitative extension of the model behind the first adaptive therapy clinical trial in metastatic prostate cancer, restated in the previous section^28^). Using the simplifying assumption assumption of a (relatively) constant tumor volume allows us to focus on the frequency dynamics within the tumor, ignoring population dynamics. Tracking frequency dynamics that are themselves frequency-dependent allows us to use a game theoretic modeling framework.

We can use the 3 by 3 payoff matrix that describes the outcomes of interactions between the different cell types. The expected payoff to an individual of a given cell type is influenced by the frequency of cell types in the population (equation 1). This can be thought of as the “inner game”^59^. The replicator equation from game theory, then translates these strategy-specific payoffs into the evolutionary dynamics described by changes in the frequencies of each cell type within the tumor (equation 2). This is the “outer game” where payoffs become translated into fitness.

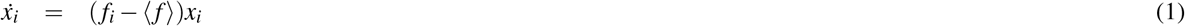

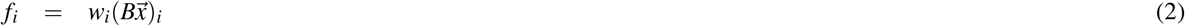

where B = 1 - A, noted above. The variables *x*_1_, *x*_2_, *x*_3_ are the corresponding frequency of dependent (*T*^+^), producers (*T^P^*) and independent (*T*^−^) cells, respectively, such that 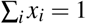. The prevalence of each cell type changes over time according to the changing payoffs, *f_i_*, compared to the average payoff of all three populations 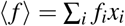. Unless otherwise noted parameters for untreated dynamics are: *b*_1,2_ = 0.2; *b*_1,3_ = 0.6; *b*_2,1_ = 0.3; *b*_2,3_ = 0.5; *b*_3,1_ = 0.4; *b*_3,2_ = 0.1, for Lupron only are: *b*_1,2_ = 0.4; *b*_1,3_ = 0.3; *b*_2,1_ = 0.6; *b*_2,3_ = 0.5; *b*_3,1_ = 0.2; *b*_3,2_ = 0.1, and for Lupron & Abirateron are: *b*_1,2_ = 0.5; *b*_1,3_ = 0.1; *b*_2,1_ = 0.6; *b*_2,3_ = 0.2; *b*_3,1_ = 0.4; *b*_3,2_ = 0.3.

The expected payoff to a cell type is calculated as the product of the payoff entries and prevalence of each population (eqn. 2). Treatments are assumed to alter the carrying capacity for each cell type. Under Lupron, each producer cell is assumed to support 1.5 dependent cells, limiting the *T*^+^ to rely on producers in the absence of systemic testosterone. During Lupron & Abiraterone treatment, each producer is capable of supporting 0.5 *T*^+^ cells and the carrying capacity of *T^P^* cells significantly drops due to local anti-androgen effects.

In order to simulate therapy, we introduced a weighting term, *w_i_*. The weighting term adjusts payoffs to a cell type by its carrying capacity (capacity of the tumor environment to support a given cell type). each cell type’s weighting term equals a given cell type’s carrying capacity, *K_i_*, normalized by a maximum carrying capacity (*w_i_* = *K_i_*/*K*_max_. Figures 4, 5, 6, 7, use the following parameters: no treatment: *K*_1_ = 1.5 · 10^4^, *K*_2_ = 10^4^, *K*_3_ = 10^2^; Lupron: *K*_1_ = 1.5 · 10^4^; *K*_2_ = 10^4^; *K*_3_ = 10^4^; Lupron & Abiraterone: *K*_1_ = 0.5 · 10^4^; *K*_2_ = 10^2^; *K*_3_ = 10^4^; *K*_max_ = 1.5 · 10^4^ cells^28^.

### Patient data model fitting

Each subpopulation is assumed to contribute to changes in prostate-serum-antigen (PSA) levels, as follows.

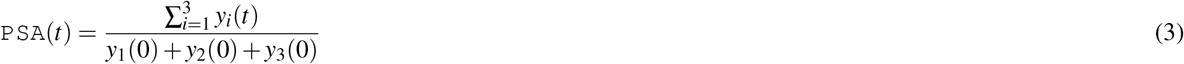

The PSA is normalized over time to show changes relative to treatment initialization (see figure 3). Best fits were performed using **lsqcurvefit** function in MATLAB 2018a. All parameters were held constant between patients (see Supplemental information) except for the initial conditions, *y_i_*(*t* = 0).

## Data and Code Availability

The data and computer code that support the findings of this study are available from the corresponding author upon reasonable request.

## Acknowledgments

The authors gratefully acknowledge funding from the Physical Sciences Oncology Network (PSON) at the National Cancer Institute, U54CA193489 (supporting J. West, J. Brown and A. Anderson) and European Union’s Horizon 2020 research and innovation program, under the Marie Skłodowska-Curie grant agreement No. 690817 (L. You).

## 1 Supplementary Information

### Population dynamics

Briefly, we restate a previously published Lotka-Volterra mathematical model used to characterize adaptive treatment of metastatic castrate resistant prostate cancer in figure 3. The model desribes competition among three cell types: testosteronedependent (T+), testosterone-producing (TP), and testosterone-independent (T−), *i* = 1,2,3, respectively.

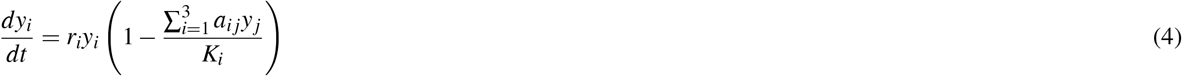

where the intrinsic growth rates, *r_i_*, were parameterized from in vitro cell data (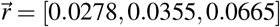; see^28^). Carrying capacities represent the contribution of each subpopulation to changes in PSA, not the overall cell count of each subpopulation. Under Lupron treatment only, the carrying capacities are given by: *K*_1_ = 1.5 · 10^4^; *K*_2_ = 10^4^; *K*_3_ = 10^4^. Lupron & Abiraterone carrying capacities are given by: *K*_1_ = 0.5 · 10^4^; *K*_2_ = 10^2^; *K*_3_ = 10^4^^28^. Carrying capacities for dynamics under no treatment are given by: *K*_1_ = 1.5 · 10^4^, *K*_2_ = 10^4^, *K*_3_ = 10^2^. Competitive interactions between these three cell types are summarized in the payoff matrix, *A*, below.

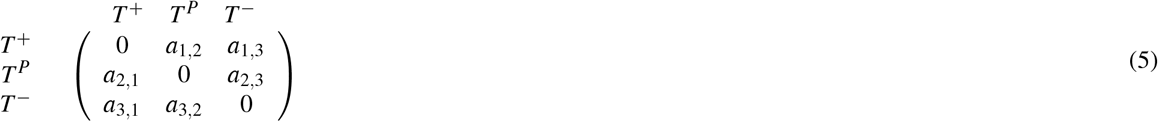

Competition parameters are described by certain inequalities for given treatment dynamics. We summarize these inequalities in Supplemental Materials for convenience: Table 1 (under treatment) and Table 2 (no treatment)^28, 45, 60^. Since there is cell-cell competition and niche partitioning with respect to association with surrounding vasculature, we assume that all coefficients are positive and bounded between 0 and 1. Consistent with ref.^28^, two general rules are used to determine the relative values of inter-cell type interactions: 1) *T*^+^ cells with no exogenous testosterone are the least competitive cell type and 2) the competitive effect of *T* cells is stronger on TP cells than on T+ cells. Unless otherwise noted parameters for untreated dynamics are: *a*_1,2_ = 0.8; *a*_1,3_ = 0.4; *a*_2,1_ = 0.7; *a*_2,3_ = 0.5; *a*_3,1_ = 0.6; *a*_3,2_ = 0.9, for Luprononlyare: *a*_1,2_ = 0.6; *a*_1,3_ = 0.7; *a*_2,1_ = 0.4; *a*_2,3_ = 0.5; *a*_3,1_ = 0.8; *a*_3,2_ = 0.9, and for Lupron & Abirateron are: *a*_1,2_ = 0.5; *a*_1,3_ = 0.9; *a*_2,1_ = 0.4; *a*_2,3_ = 0.8; *a*_3,1_ = 0.6; *a*_3,2_ = 0.7.

